# Natural methylation epialleles correlate with gene expression in maize

**DOI:** 10.1101/2023.01.23.525249

**Authors:** Yibing Zeng, R. Kelly Dawe, Jonathan I. Gent

## Abstract

DNA methylation (5-methylcytosine) represses transposon activity and contributes to inaccessible chromatin structure of repetitive DNA in plants. It is depleted from cis regulatory elements in and near genes but is present in some gene bodies, including exons. Methylation in exons solely in the CG context is called gene body methylation (gbM). Methylation in exons in both CG and non-CG contexts is called TE-like methylation (teM). Assigning functions to both forms of methylation in genes has proven to be challenging. Toward that end, we utilized recent genome assemblies, gene annotations, transcription data, and methylome data to quantify common patterns of gene methylation and their relations to gene expression in maize. To compare between genomes, we analyzed each data source relative to its own genome assembly rather than the easier but less accurate method of using one assembly as reference for all. We found that gbM genes exist in a continuum of CG methylation levels without a clear demarcation between unmethylated genes and gbM genes. Analysis of expression levels across diverse maize stocks and tissues revealed a weak but highly significant positive correlation between gbM and gene expression except in endosperm. gbM epialleles were associated with an approximately 3% increase in steady-state expression level relative to unmethylated epialleles. In contrast to gbM genes, which were conserved and were broadly expressed across tissues, we found that teM genes, which make up about 12% of genes, are mainly silent, are limited to specific maize stocks, and exhibit evidence of annotation errors. We used these data to flag all teM genes in the 26 NAM founder genome assemblies. While some teM genes are likely functional, these data suggest that the majority are not, and their inclusion can confound interpretation of whole-genome studies.

## INTRODUCTION

DNA methylation is one part of a multilayered chromatin-based method of repressing transcription and accessibility of repetitive DNA in plants. The mode of repression depends in part on the two or three nucleotide sequence context of the methylated cytosine, generally categorized as CG, CHG, and CHH, where H = A, T, or C. DNA methylation is not restricted to repetitive DNA, however. In most flowering plants, a large fraction of genes can have methylation in the CG context (mCG) in exons [reviewed in (Bewick and Schmitz 2017; Muyle *et al*. 2022)]. Such genes are referred to as gene body methylated genes, gbM genes for short. Genes that have TE-like methylation, both mCG and non-mCG, in their exons are referred to as teM genes. The third and most abundant group of genes are unmethylated in their exons and referred to as UM genes. All three methylation groups have signature expression patterns: gbM genes tend to be broadly expressed across tissues, UM genes tend to be tissue-specific, and teM genes tend to be poorly expressed [reviewed in (Bewick and Schmitz 2017; Muyle *et al*. 2022)].

The mCG in gbM genes is maintained by methyltransferases of the MET1 family (Stroud *et al*. 2013). However, there is no dedicated mechanism to establish gbM on genes where it is absent: Genes that have lost gbM in *met1* mutants do not reacquire gbM after MET1 is returned (at least over a period of eight generations (Reinders *et al*. 2009). Instead, establishment of mCG in gbM genes is hypothesized to occur slowly and infrequently through intermediate teM-like states mediated by spurious DNA methylation by MET1 and chromomethyltransferases of the CMT family (Niederhuth *et al*. 2016; Wendte *et al*. 2019). Although the direct output of CMTs is mCHH or mCHG, they can also lead to mCG by downstream recruitment of MET1 in Arabidopsis (Lyons *et al*. 2022). According to this hypothesis, histone demethylation by the Jumonji C histone demethylase leads to loss of CMT activity and associated mCHG (Saze *et al*. 2008). Continued activity of MET1 coupled to DNA replication causes mCG to persist, leading to an epigenetically stable gbM state. Supporting this view are data showing that CMTs can target gbM genes in Arabidopsis (Zhang *et al*. 2020, 2021; Papareddy *et al*. 2021). Expression of Arabidopsis CMT in *Eutrema salsugineum* (which normally lacks both CMT and gbM) can induce mCG in gene bodies, which persists after CMT is removed (Wendte *et al*. 2019).

Since cytosine methylation increases the frequency of G:C to A:T transitions (Ossowski *et al*. 2010), mCG in coding DNA might be harmful rather than beneficial, unless it provides other benefits that outweigh its mutagenic tendency. Gene body methylation is absent from the vast majority of fungal genomes, even in genomes with methylation at repetitive elements (Bewick *et al*. 2019). The existence of gbM in diverse animals, however, is evidence that beneficial functions likely exist. In fact, some insect genomes have extensive gene body methylation but little methylation of repetitive elements (Bewick and Schmitz 2017). Vertebrates have gene body methylation as well as a dedicated mechanism for establishing it: recruitment of DNMT3 methyltransferases coupled to trimethylation of histone H3 lysine 36 (H3K36me3) (Baubec *et al*. 2015; Bröhm *et al*. 2022). In plants, H3K36me3 seems to be unrelated to gbM (Wollmann *et al*. 2017). The functional significance of gbM in vertebrates is unclear but may prevent internal transcriptional initiation (Neri *et al*. 2017; Teissandier and Bourc’his 2017). In some cases, gbM also impacts splicing (Yearim *et al*. 2015; Shayevitch *et al*. 2018). Function of gbM could be compared to the repression of transposons and other repetitive elements by DNA methylation: Although they are strongly methylated in some eukaryotes, they are poorly methylated or not methylated at all in others, including nematodes, fruit flies, yeasts, and honeybees [reviewed in (Schmitz *et al*. 2019)]. Another appropriate comparison might be the centromeric histone variant CENP-A, which is essential in most eukaryotes but absent in some (Drinnenberg *et al*. 2014).

Whether mCG in gbM genes has a biologically significant effect on steady-state mRNA levels in plants is not clear. Comparisons of gene expression in *E. salsugineum* (without gbM) with *A. thaliana* (with gbM) has yielded conflicting evidence both for and against gbM promoting gene expression (Muyle and Gaut 2019; Bewick *et al*. 2019). Experiments using methylation data from the Arabidopsis 1001 Genome Consortium (Kawakatsu *et al*. 2016) found evidence of selection on gbM epialleles as well as small increases in expression in gbM over UM epialleles (Shahzad *et al*.; Muyle *et al*. 2021). Older studies based on other sets of Arabidopsis accessions also revealed small but positive correlations between gbM and gene expression levels (Schmitz *et al*. 2013; Dubin *et al*. 2015; Meng *et al*. 2016). Expression analysis of genes that lost gbM through *met1* mutation in Arabidopsis in theory should provide the most direct evidence for a causal effect of gbM on gene expression. However, such studies have produced conflicting results both for and against gbM promoting gene expression (Shahzad *et al*.; Bewick *et al*. 2019). The fact that gbM genes tend to be expressed more broadly through development than UM genes raises the possibility that gbM may function in stabilizing gene expression across development, with subtle activating (or repressing) functions depending on the cell type (Takuno and Gaut 2012, 2013; Niederhuth *et al*. 2016). It is also possible that gbM has a function unrelated to normal gene regulation–for example, inhibiting ectopic initiation of transcription in gene bodies. The evidence for such a function is mixed in plants (Choi *et al*. 2020; Le *et al*. 2020). One might also speculate functions for gbM unrelated to transcription.

DNA glycosylases, which demethylate DNA through the base excision repair pathway, provide strong evidence that DNA methylation can function in gene regulation. For example, DNA glycosylases can function to activate genes upon bacterial infection (Halter *et al*. 2021), in response to abscisic acid hormone signaling (Kim *et al*. 2019), and in pollen tube development (Khouider *et al*. 2021). In endosperm, DNA glycosylases act on maternal alleles of some genes to cause genomic imprinting, which is essential for endosperm development [reviewed in (Anderson and Springer 2018)]. In the microgametophyte, including mature pollen, demethylation by DNA glycosylases has been proposed to take on a larger role in gene regulation than in sporophytic cells (Borg *et al*. 2021). While TE-like methylation of TEs themselves is generally stable across sporophytic development (Crisp *et al*. 2020), TE-like DNA methylation in coding DNA might indicate a function in gene regulation.

Similar associations between gene body methylation with broad gene expression patterns, conservation, and gene structural features in Arabidopsis also hold true in other plants (Takuno and Gaut 2013; Niederhuth *et al*. 2016; Seymour and Gaut 2020; Martin *et al*. 2021b). Whether the positive correlation between gbM epialleles and gene expression holds true in other plants has not been rigorously tested. Maize in particular provides an interesting test case because of its different genome and epigenome structure, including DNA methylation patterns and much higher repeat content near genes (Gent *et al*. 2013; Hufford *et al*. 2021). In addition, its genome contains two subgenomes, which allows for large changes in methylation and expression of many genes to be compensated for by unchanged, second-subgenome copies (Woodhouse *et al*. 2010). The promoters of most functional maize genes are constitutively demethylated, and methylation in promoters is a strong indicator of silencing (Hufford *et al*. 2021). Recently, improved genome assemblies and annotations were produced for the B73 inbred line and 25 other diverse inbred lines known as the NAM founders (Hufford *et al*. 2021). Transcriptome data for ten tissues and 20X coverage methylome data for developing leaves for each genome provide an opportunity to better characterize gene methylation trends in maize. We made use of this resource to identify natural epialleles and explore the relationships between methylation (UM, gbM, and teM) and gene expression on a pan-genome scale.

## RESULTS

We first surveyed the landscape of gene methylation in maize using the B73 genome and a DNA methylome from developing seedling leaves as a reference (Hufford *et al*. 2021). Introns often have TE-like methylation that is distinct from flanking exons simply because they contain TE insertions (Seymour and Gaut 2020). The very 5’ and 3’ end of UTRs are typically unmethylated when they are correctly annotated, but UTR annotations are often imprecise and sometimes overlap with nearby TEs. Thus we excluded both UTRs and introns in measuring genic methylation. For each gene with sufficient read coverage (reads spanning at least 40 CGs and 40 CHGs), we assigned a single mCG value and mCHG value as the average methylation in its coding DNA sequence (CDS). These values are measured as a proportion of methylated cytosines to total cytosines, and range from zero to one. To produce a visual summary of gene methylation trends, we represented each gene by these two values. A clear bimodal trend was evident, where the larger group of genes had low mCHG methylation and a continuous range of mCG but heavily skewed toward zero mCG (Fig. 1A). A second, smaller group of genes had both high mCG and mCHG (Fig. 1A). We divided the first group into UM genes and gbM genes, where UM had less than or equal to 0.05 mCG and gbM had at least 0.2 mCG. Both UM and gbM had less than or equal to 0.05 mCHG. We defined genes in the second group as teM genes based on at least 0.4 mCG and 0.4 mCHG. This produced 14,393 UM genes, 8,134 gbM genes and 3,402 teM genes. The remaining 4,277 genes with intermediate methylation values were left uncategorized, along with 9,550 genes with insufficient methylome sequencing read coverage to confidently assign methylation epiallele status.

**Figure 1.**
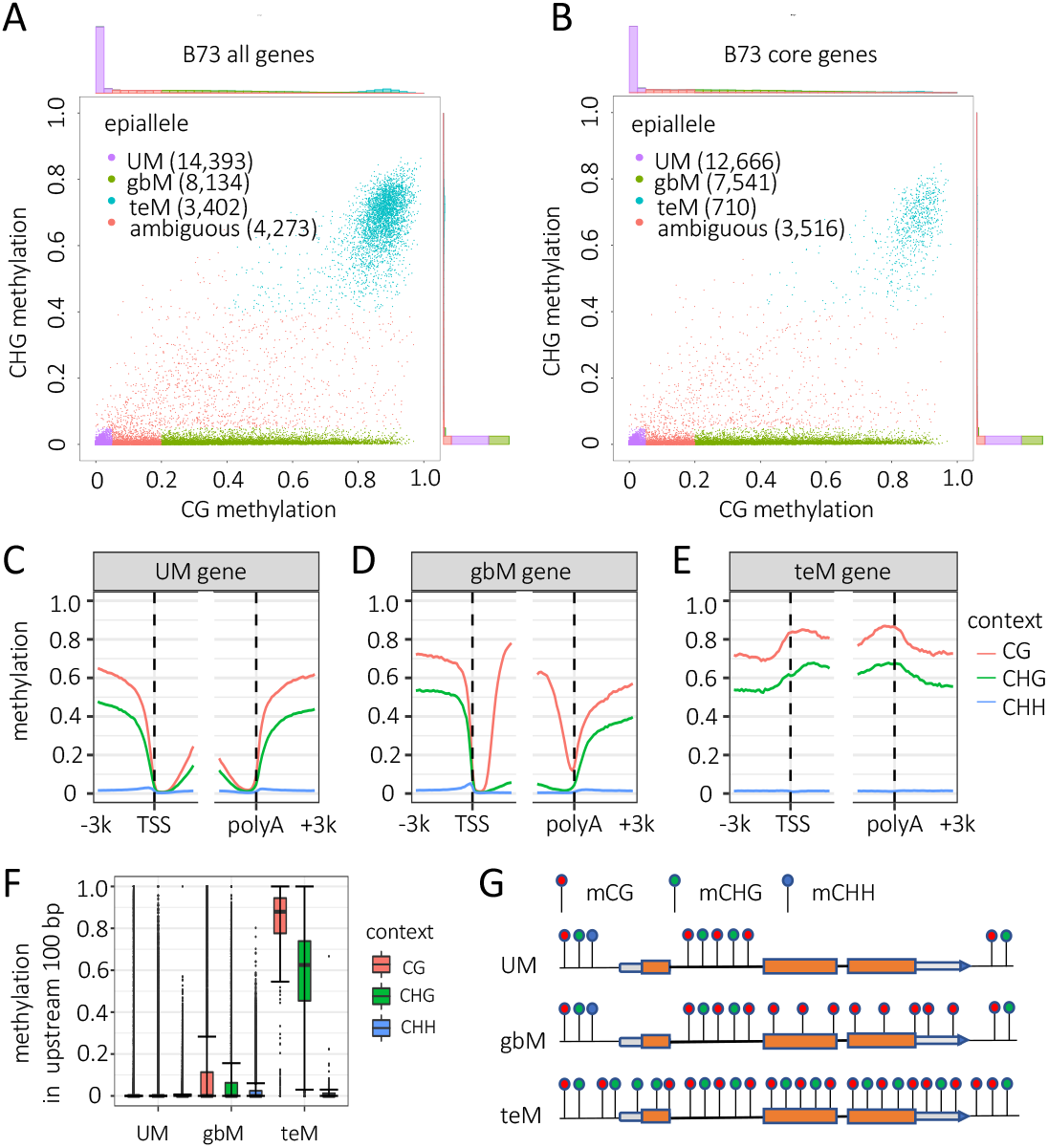
**A-B**. Scatter plots of mCG vs. mCHG for all B73 genes (A) and core B73 genes (B). These values are measured as a proportion of methylated cytosines to total cytosines, and range from zero to one. Values indicate methylation of coding DNA sequence (CDS) only. Histograms outside axes indicate gene counts in each range of methylation values. Only genes with sufficient coverage of EM-seq reads were included in this analysis (at least 40 cytosines in each context spanned by reads). **C-E**. Metagene mCHH, mCHG, and mCG for core UM, gbM, teM B73 genes. Genes were aligned at transcription starts sites (TSSs) and polyadenylation sites (polyA). The plots show 3 Kb upstream, 3 Kb downstream, and 2 Kb of internal sequence. Methylation values are measured in 100-bp intervals. **F**. Distribution of mCHH, mCHG, and mCG for the upstream 100-bp regions for UM, gbM, and teM B73 core genes. Whiskers indicate 1.5 interquartile range (IQR). **G**. Schematic of gene methylation (epiallele) types. Lollipops indicate methylated cytosines, color coded by context.

To enrich for functional genes over gene annotation artifacts, we made use of the core gene categorization scheme previously developed by comparison of the 26 NAM founder genomes (Hufford *et al*. 2021). Core genes are the subset that are present at syntenic positions in all 26 genomes, based on sequence homology. These include all copies of tandemly duplicated genes and gene fragments. Core genes are enriched for synteny with Sorghum and for detectable RNA expression relative to the complete set of annotated genes (Hufford *et al*. 2021). 28,292 of 39,756 annotated B73 genes are core genes. Repeating the above gene methylation categorization scheme with only core genes had little effect on the numbers of UM and gbM genes and retained the continuum of mCG in these categories, but it produced a 78% decrease in the proportion of teM genes, down from 3,402 to 710 (Fig. 1B). This decrease in the number of teM genes among core genes suggests that many are pseudogenes or mis-annotated TEs. For all subsequent analyses, we included only core genes, unless otherwise indicated. While we defined teM genes solely based on methylation in CDS, they also had high mCG and mCHG at their annotated transcription start sites (TSSs) and polyadenylation sites.

This was evident both from metagene methylation profiles (Fig. 1C-E), as well as the distribution of methylation levels in the first 100 bp upstream of TSSs (Fig. 1 F). mCHH upstream of TSSs is a common feature of genes (Gent *et al*. 2013; Martin *et al*. 2021a). In contrast to UM and gbM genes, teM genes showed no enrichment for mCHH upstream of genes (Fig. 1C-E). Figure 1G provides a simplified summary of methylation profiles for the three gene methylation types.

Consistent with longer lengths of Arabidopsis gbM genes (Zhang *et al*. 2006; Zilberman *et al*. 2007; Takuno and Gaut 2012), comparison of structural features of the genes in each methylation category revealed that all components of maize gbM genes (UTRs, CDSs, introns, and intronic TEs) were longer than UM genes, producing an average 2.9-fold longer total gene lengths (Fig. 2A). Differences were significant for all features (p-value < 10^−9^, two-tailed Wilcoxen rank sum test). The total-gene length differences were at least partly explained by the number of exons and introns: The mean number of exons was 3.2 for UM genes, 9.8 for gbM, and 3.4 for teM (Supplemental Fig. 1). Introns had the largest difference in length: the average cumulative intron length of gbM genes was 4,437 bp longer than UM genes (5,481 bp - 1,144 bp). 1,576 bp of this difference in length could be accounted for by TEs in introns (1,965 bp in gbM, 390 in UM average cumulative lengths of TEs in introns). Introns of gbM genes were more likely to contain TEs of all superfamilies (Fig. 2B). teM genes were also distinguished from both gbM and UM genes by short or absent UTRs (Fig. 2A and C). 59.1% of teM genes lacked both 5’ and 3’ UTRs, compared to 0.38% of gbM and 6.3% of UM genes. teM genes had relatively short CDSs (816 bp for teM genes, compared to 1,858 bp for gbM and 1,079 bp for UM genes) and tended to overlap annotated TEs: 34.6% of teM genes had at least 100 bp of overlap between CDS and TEs, compared to 5.3% of gbM and 3.4% of UM genes (Fig. 2D).

**Figure 2.**
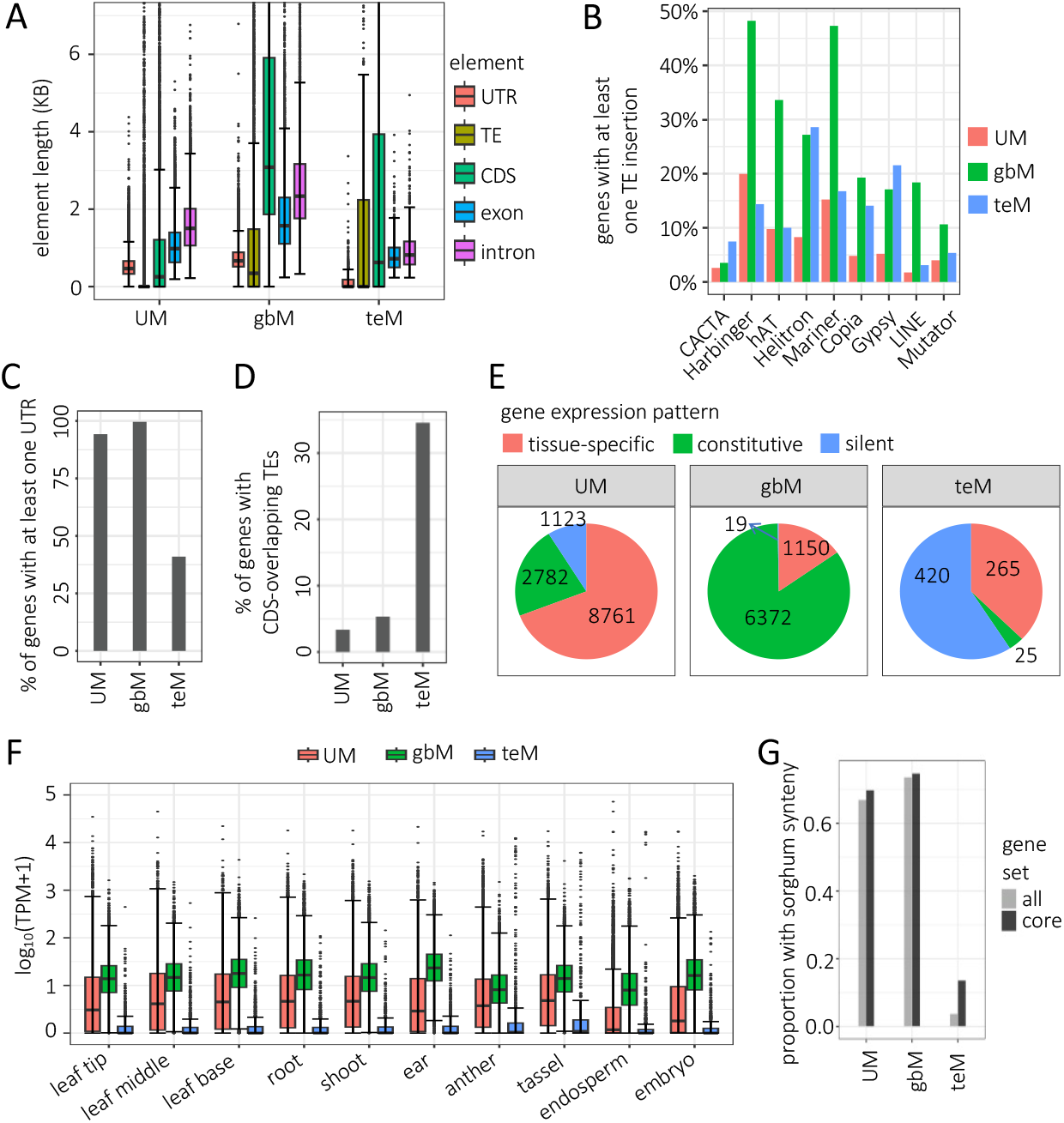
**A**. Distributions (left to right) of total CDS length, exon length, intron length, intron-overlapping TE length, and UTR length in UM, gbM and teM genes. Lengths are cumulative such that individual elements in a gene are summed to yield a single value for the gene. Whiskers indicate 1.5 IQR. Length differences between UM, gbM and teM genes were significant for all features (p-value < 10^−9^, two-tailed Wilcoxen rank sum test). **B**. Proportion of genes containing at least one TE insertion overlapping introns. All nine TE superfamilies are significantly different between UM and gbM (P-value < 0.01, one-tailed Mann-Whitney test). **C**. Proportion of genes with at least one annotated UTR. **D**. Proportion of genes with at least 100 bp overlap of CDS and annotated TEs. **E**. Proportion and number of genes in each gene expression category. Each pie represents all genes with a defined epiallele type (UM, gbM, and teM), with the number of genes in each expression category indicated for each slice. These are arranged clockwise starting with tissue-specific expression at 12 o’clock, then constitutive expression, then silent. **F**. Distribution of TPM values for UM, gbM and teM genes across tissue types. Log base 10 of TPM values (plus a pseudocount of 1 to avoid problems with zero values) are shown, and Y-axis is truncated at +5 and -5. **G**. Percent of genes with syntenic homologs in *Sorghum bicolor*. The differences between each category (UM gbM, and teM) were significant for all genes and core genes (P-value < 10^−10^, two sample Z proportion test).

To investigate expression patterns for each category of gene methylation, we used RNA-seq data from ten tissues for each NAM founder inbred (Hufford *et al*. 2021). UM genes had larger expression ranges in each tissue than gbM genes, showing both higher and lower extremes in TPM values, consistent with tissue-specific expression (Fig. 2E, F). In contrast, gbM genes were consistently expressed at moderate levels across the ten tissues, consistent with constitutive gene expression (Takuno and Gaut 2012, 2013; Niederhuth *et al*. 2016). This moderate expression across tissues results in an average higher expression of gbM genes than UM genes (Fig. 2F). In ear for example, the average gbM expression was 40.7 TPM and UM was 25.7 TPM. teM genes were poorly expressed across all tissues. Anther, tassel, and endosperm had a higher skew of teM TPM values than the other seven tissues, suggesting that a number of teM genes are highly expressed in these tissues. We categorized genes as tissue-specific based on a TPM value of less than one in at least one but not all tissues, as constitutively expressed based on a TPM value of at least one in all ten tissues, or as constitutively silent based on a TPM value of less than one in all tissues. This expression categorization scheme correlated with methylation categories, where UM genes were most tissue-specific, gbM most constitutive, and teM most silent (Fig. 2E).

Even though we only included core genes in these analyses, the poor expression of teM genes, frequent absence of UTRs, and high overlap with TEs raised the question of whether they are conserved outside of maize. To test this, we asked how many had unambiguous homologs at syntenic positions in sorghum and could be assigned to one of the two maize subgenomes (Woodhouse *et al*. 2010; Hufford *et al*. 2021). While 75% of gbM genes and 70% of UM genes had syntenic homologs, only 14% of teM genes did (Fig. 2G). When including the entire set of B73 genes rather than just core genes, 74% of gbM genes and 67% of UM genes had syntenic homologs, but only 4% of teM genes did.

We applied the same methods used in B73 to categorize genes as UM, gbM or teM to the 25 other NAM founder genomes. Specifically, we used the methylome data previously analyzed, where each set of EM-seq reads was mapped to its own genome and analyzed with respect to its own gene annotations (Hufford *et al*. 2021). For this analysis we included all gene annotations, not just core genes. The abundance of identified UM genes varied from 51% of total genes annotations with sufficient EM-seq read coverage in CML247 to 44% in M37W (Supplemental Fig. 2). The abundance of gbM genes varied from 18% in CML247 to 28% in M37W. The higher abundance of UM genes and lower of gbM genes in CML247 is consistent with the low genome-wide mCG previously observed in CML247 (Hufford *et al*. 2021). The abundance of teM genes hardly varied, from 11% in Ky21, to 13% in Ki3. Although some teM genes may be demethylated as a means to regulate gene expression, most are likely mis-annotations or pseudogenes. The teM genes are displayed as genome browser tracks for all 26 genomes hosted by the Maize Genetics and Genomics Database (MaizeGDB) (Woodhouse *et al*. 2021). For examples, see Supplemental Figure. 3. The teM genes are also listed along with UM and gbM genes for all 26 genomes at https://github.com/dawelab/Natural-methylation-epialleles-correlate-with-gene-expression-in-maize/tree/main/patterns%20of%20gene%20methylation/epiallele.

When genes are duplicated, their methylation or expression states may change. To specifically test for epiallele switches associated with tandem duplication, we looked for genes that were present as singletons in at least two genomes but as tandem duplicates in at least one other. For this purpose, we used the pangene system to link genes across genomes (Hufford *et al*. 2021). All genes that have homologous sequence at a syntenic position in the genomes share the same pangene name. For all analyses, we included only the subset of 27,910 of the 103,033 pangenes which are core pangenes (pangenes with genes in all 26 genomes). Some core pangenes have more than 26 genes because multiple tandem duplicate genes are linked to single pangenes (Fig. 3A). However, tandem duplications often capture only a fragment of the gene. Also, a single gene can also appear to be a tandem duplicate because 5’ and 3’ portions are incorrectly annotated as separate genes. To avoid both these issues, we included only intact tandem duplicates for all pangene analyses. We defined intact tandem duplicates as those whose cumulative CDS lengths differed by no more than 10% from the median length of all singletons for a given pangene. Since gene dosage effects provide a strong constraint on gene copy number and (Birchler and Veitia 2012), and singletons predominate over tandem duplicates in plant genomes, we assumed for the purposes of this analysis that singletons best represent ancestral epialleles and tandem duplicates best represent derived ones. 6,695 pangenes exist as tandem duplicates in some inbreds and singletons in at least two other inbreds. Among these 6,695 pangenes, 725 of the singletons were present in more than one epiallele state. We excluded these from further analysis to avoid ambiguity about which epiallele represents the singleton state. The remaining 5,970 pangenes, represented by singletons of one epiallele type in at least two genomes, were represented by arrays of up to 47 duplicates in other inbreds. We refer to these as “1-to-N” pangenes.

**Figure 3.**
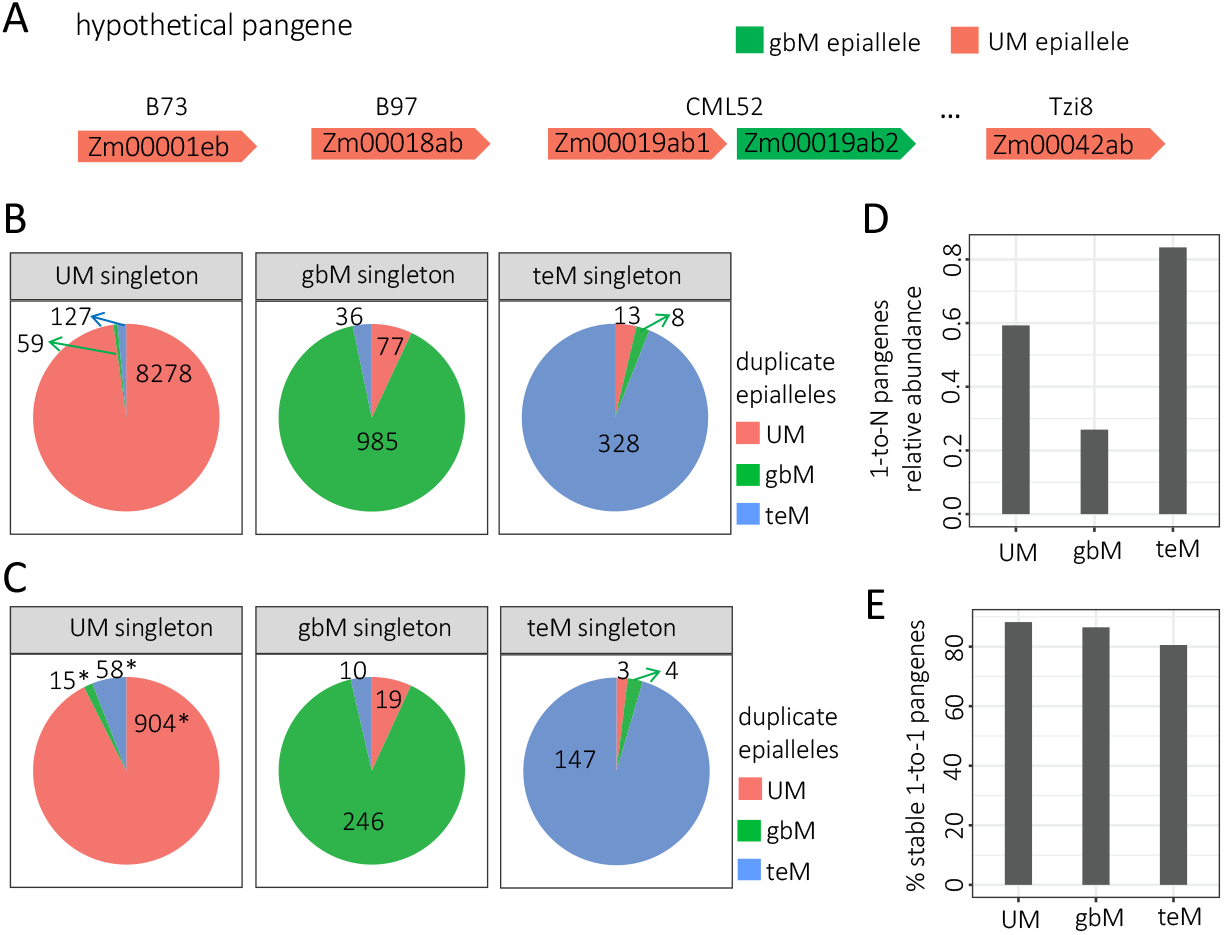
**A**. Schematic of the relationship between a hypothetical 1-to-N pangene and its individual genes. **B**. Abundance of each epiallele type among tandem duplicate genes. Each pie represents all tandem duplicates with one singleton epiallele type, and the number listed in each slice indicates the number of tandem duplicates with UM, gbM, or teM epialleles: These are arranged clockwise starting with UM epialleles at 12 o’clock. **C**. Abundance of each epiallele type among tandem duplicate genes, but only including tandem duplication genes with a copy number of at least four. Asterisks indicate a significant difference in proportion from the corresponding value in panel B, (P-value < 1 × 10^−3^, Chi-square test). **D**. Relative abundance of 1-to-N pangenes. 1-to-N pangenes were categorized by their singleton epiallele type, and the total number divided by the number of stable 1-to-1 pangenes of the same epiallele type. **E**. Percent of 1-to-1 pangenes with stable epialleles. The number of 1-to-1 stable pangenes of each epiallele type was divided by the total number of unstable and stable 1-to-1 pangenes with that epiallele.

**Table 1:** Usage of key terms

|  |  |
| --- | --- |
| UM | unmethylated in coding DNA sequence (CDS) |
| gbM | gene body methylation, only mCG in CDS |
| teM | TE-like methylation, mCG and mCHG in CDS |
| epiallele type | methylation status of genes: UM, gbM, or teM |
| tissue-specific gene | expressed in at least one tissue and not expressed in at least one other tissue |
| constitutive gene | expressed in all ten tissues examined |
| silent gene | not expressed (silent) in all ten tissues |
| NAM founders | set of 26 diverse maize inbred stocks including B73 |
| core gene | present at syntenic position in all NAM founders |
| pangene | set of homologous genes at a syntenic position in different genomes |
| 1-to-1 pangene | pangene that has one intact copy in all NAM founders |
| 1-to-N pangene | pangene with intact tandem duplicates in NAM founders |
| stable pangene | represented by only one epiallele in NAM founders |
| unstable pangene | represented by more than one epiallele in NAM founders |

Comparing the singleton epialleles with their corresponding duplicate epialleles in other genomes revealed that epiallele states of the singletons were normally maintained in the duplicates (Fig. 3B). In the case of UM singletons, 98% of duplicates were also UM. These are surprisingly high percentages because a change in mCG value as small as 0.15 is enough to switch between UM and gbM. A larger change of 0.35 in both mCG and mCHG is required for changing between UM and teM, yet switches of this type were more common than UM to gbM (1.5% teM duplicates vs 0.7% gbM duplicates for UM singletons). Abundance of teM duplicates was higher (5.9%) in the subset of cases with a minimum copy number of four (Fig. 3C). This increase in teM duplicates in the minimum-four-copy set was highly significant (P-value < 10^−10^, Chi-square test). In the case of gbM singletons, 89% of duplicates were also gbM, with 6.9% UM duplicates and 3.6% teM duplicates. These numbers were nearly identical in the set of duplicates with a minimum copy number of four.

Of the 5,970 1-to-N pangenes, 4,149 had UM singleton epialleles, 1,589 had gbM singleton epialleles, and 232 had teM singleton epialleles. To put these numbers in perspective, we compared the numbers to stable 1-to-1 pangenes, which had only singletons and only one epiallele type. As with 1-to-N pangenes, partial duplications were ignored in determining singleton vs. duplicate status. After excluding pangenes that had fewer than two genes with defined epialleles, 7,001 of the stable 1-to-1 pangenes had only UM epialleles, 5,992 had only gbM, and 277 had only teM. Normalizing the numbers of 1-to-N pangenes by the numbers of stable 1-to-1 pangenes suggests that UM singletons are 2.2-fold more likely to be associated with gene duplications than gbM singletons (Fig. 3D). gbM epialleles may be less susceptible to tandem duplication or have more severe fitness consequences than duplicates derived from UM epialleles.

An additional 964 core pangenes occurred as singletons in all genomes but were present in more than one epiallele state. We called these unstable 1-to-1 pangenes. There were 897 UM-gbM unstable 1-to-1 pangenes, 27 UM-teM, 30 gbM-teM, and 10 UM-gbM-teM. To be included in one of these unstable 1-to-1 pangene groups, we required that at least two genomes contain each epiallele. The remaining 7,706 core pangenes did not meet the stringent requirements for 1-to-N, stable 1-to-1, or unstable 1-to-1 pangenes. A total of 13% of the UM, 15% of the gbM, and 19% of the teM epiallele-containing pangenes were unstable (calculated as unstable 1-to-1 pangenes divided by stable plus unstable 1-to-1 pangenes) (Fig. 3E).

Maize is a pseudotetraploid where many genes have two copies with potentially redundant functions (Woodhouse *et al*. 2010; Hufford *et al*. 2021). Thus maize offers an opportunity to test relationships between epialleles and gene expression changes that would not be tolerated in plants where most genes are single copy. For each of the 897 unstable 1-to-1 UM-gbM pangenes, we calculated the differences in the mean transcripts per million (TPM) for UM epialleles and gbM epialleles, which we refer to as the gbM-UM TPM differences. Epialleles with large TPM values create a broad distribution of gbM-UM TPM differences. Nonetheless, the distribution of differences would be expected to center on a value of zero if UM-gbM epiallele switches are not associated with gene expression change. We found that the median gbM-UM TPM difference was above zero for all ten tissues (Figure 4A and B). These differences were significant by binomial sign tests for all tissues except endosperm (P-value < 10^−10^ except endosperm P-value = 0.07). The differences were also significant by Wilcoxon signed rank tests (P-value < 0.05 except endosperm P-value = 0.16). Normalizing the median gbM-UM TPM differences by mean UM TPM values for the same set of pangenes in each tissue produced highly consistent results. The median gbM-UM TPM differences indicate that gbM epialleles are expressed about 3% higher than UM epialleles. The differences were lowest in endosperm (at 0.5%) and highest in leaf tip (at 4.3%). We used median values for these analyses because the means are skewed by epialleles with large TPM values, where very minor changes in expression manifest as very large changes in TPM value. For example, a 10% change in expression of a gene with a TPM value of 1000 would give a gbM-UM TPM difference of 100 TPM. Even so, all tissues but root yielded mean gbM-UM TPM difference of greater than zero. As comparisons, we also examined epiallele expression in the 27 UM-teM and 30 gbM-teM unstable 1-to-1 pangenes. The small numbers of these pangenes preclude meaningful quantification of expression changes for teM epialleles, but there was a clear trend for teM epialleles to have reduced expression relative to UM or gbM (Fig. 4C and D).

**Figure 4.**
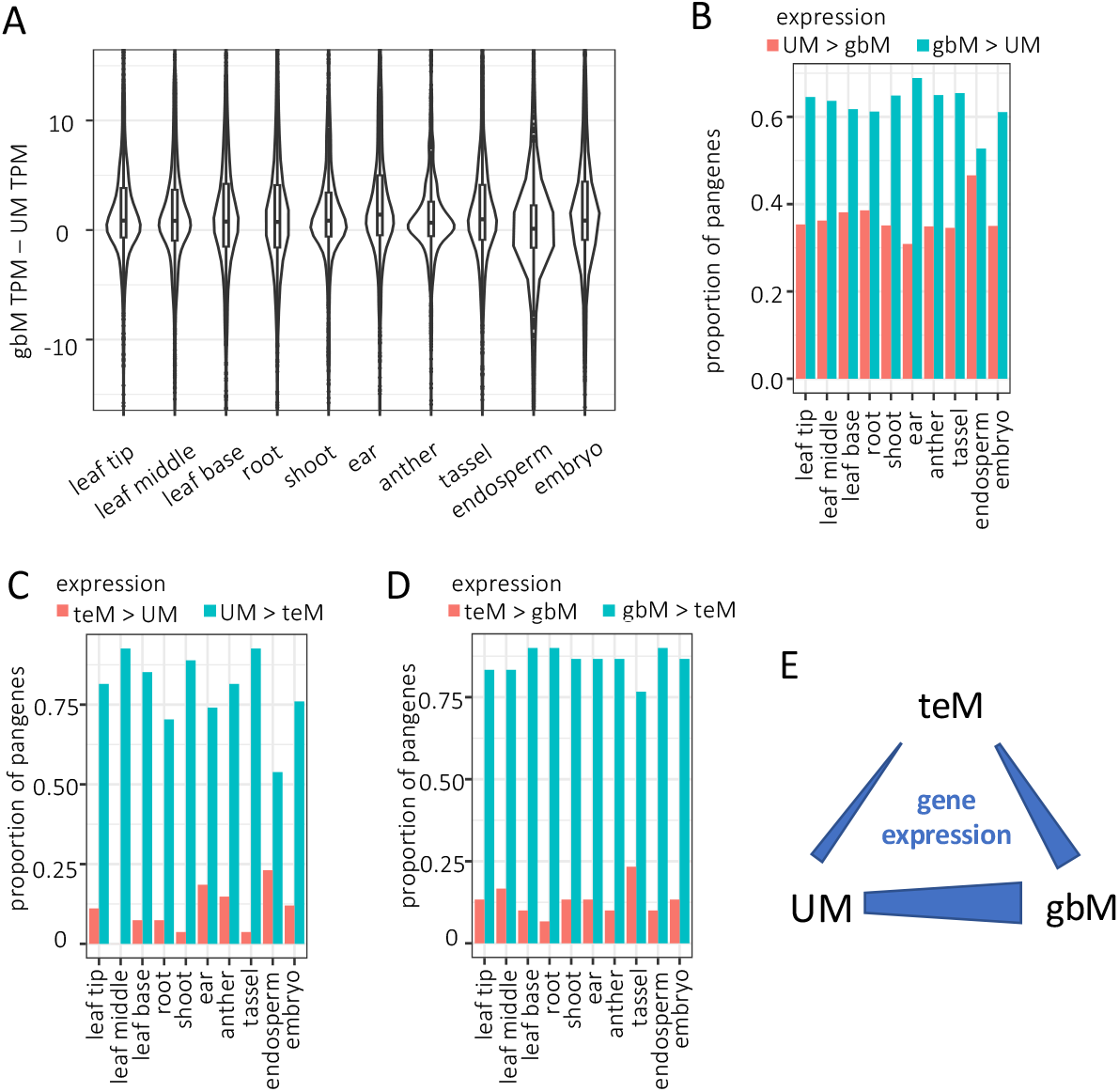
**A**. Distribution of differences in TPM between gbM and UM epialleles for each 1-to-1 unstable pangene. **B**. Proportion of 1-to-1 unstable pangenes with differences in TPM between gbM and UM epialleles that were either greater or less than zero. **C**. Proportion of 1-to-1 unstable pangenes with differences in TPM between UM and teM epialleles that were either greater or less than zero. **D**. Proportion of 1-to-1 unstable pangenes with differences in TPM between gbM and teM epialleles that were either greater or less than zero. **E**. Schematic of relationship between epialleles and gene expression.

As an alternative means of testing for a correlation between gbM and expression, we treated mCG as a quantitative variable rather than using the binary system of UM vs. gbM. This also allowed us to include genes with intermediate mCG levels (between 0.05 and 0.2). All other criteria were unchanged, such as exclusion of genes with mCHG levels above 0.05 and requirement for read coverage of at least 40 CGs and 40 CHGs per gene. As there are limitless factors that could in theory contribute to gene expression changes, of which mCG is likely to be a very minor one, we only included the potentially informative pangenes with variation in mCG. Specifically, we required that all pangenes have at least one pair of genes whose mCG difference exceeded 0.2. For example, if the gene with the lowest mCG value was 0.1, another gene would need to have a value greater than 0.3 for that pangene to be included in the analyses. In total, 8547 pangenes met this criteria. We then calculated correlation coefficients between mCG and TPM for each pangene. While we expected that correlation coefficients for individual pangenes would vary hugely, the distribution of values would reveal whether there was an overall positive trend. Indeed, both the median and mean correlation coefficients were positive for nine of the ten tissues (Supplemental Figure 4A, B). The interesting exception was endosperm, a tissue which is known to be epigenetically distinct from other tissues examined, including maternal DNA demethylation at thousands of loci (Batista and Köhler 2020; Gent *et al*. 2022). We also carried out a parallel analyses using mCHG as a quantitative trait instead of mCG to compare with TPM values. In this analysis, we entirely ignored mCG, but we retained the requirement for at least 40 CGs per gene to ensure a similar starting set of genes. 927 pangenes met the required criteria, including having at least one pair of genes whose mCHG difference exceeded 0.2. In all tissues, the ratio of positive to negative correlation coefficient values was less than one (Supplemental Figure 4C, D). These results confirm the positive correlation between gbM and expression as well as the negative correlation between teM and expression.

## DISCUSSION

### Gene body methylation defines a continuum of genes, while TE-like methylation defines a distinct group that is enriched for annotation artifacts

Examining CG methylation alone reveals a clear bimodal distribution, with most genes having either near-zero CG methylation or greater than 70% methylation (Fig.1A and B). After removing the set of genes with high CHG methylation, which is characteristic of heterochromatin, the remaining genes show a range of CG methylation heavily weighted toward zero CG methylation. Thus a categorization of non-CHG methylated genes as either unmethylated (UM) or gene body methylated (gbM) is a simplification of a continuously distributed feature. Despite the limitations of simplified UM-gbM binary categorization schemes, they do correlate with structural and expression features and may reveal hints of both the origins and consequences of gene body methylation [Reviewed in (Bewick and Schmitz 2017; Muyle *et al*. 2022)].

Examining both CG and CHG methylation together, however, reveals a distinct TE-like methylation (teM) category. We found multiple lines of evidence that this category is dominated by nonfunctional genes and annotation artifacts. First, they are poorly conserved among maize lines, with 79% of the teM gene annotations in B73 being absent from at least one of the other 25 genomes (Fig. 1A, B). Even more strikingly, only 4% had syntenic homologs in sorghum (Fig. 2G). Second, their methylation extends into their 5’ and 3’ flanking sequences, suggesting a lack of functional cis regulatory sequence (Fig. 1C, D). Third, they frequently lack both 5’ and 3’ UTRs, have short CDSs, and their CDSs overlap with annotated TEs (Fig. 2A, C, D). Fourth, they are very poorly expressed, even the 21% of teM genes in B73 that were annotated as genes in all 26 NAM founders (Fig. 2E, F and 4C, D). These data strongly suggest that most teM genes are nonfunctional. They are likely pseudogenes, fragmented tandem duplications, TEs, or other annotation errors.

Multiple lines of evidence suggests that keeping gene regulatory elements constitutively free of TE-like methylation keeps them competent for activation, even when they are currently in repressed states. In some cases where TEs have inserted into regulatory elements, DNA methylation can result in developmentally responsive repression of functional genes. FLOWERING WAGENINGEN (FWA) and REPRESSOR OF SILENCING 1 (ROS1) are well-studied examples in Arabidopsis (Fujimoto *et al*. 2008; Williams *et al*. 2015). Some functional genes also utilize TE-like methylation as a means of developmental gene regulation independently of TEs, as demonstrated by several genomically imprinted genes. For example MEDEA (MEA) and FERTILIZATION INDEPENDENT ENDOSPERM2 (FIE2), are both activated in endosperm by DNA demethylation in Arabidopsis (Batista and Köhler 2020). FWA is also in this category, as are a small number of genes in pollen (Khouider *et al*. 2021; Borg *et al*. 2021). We expect that the subset of teM genes that are present in all 26 genomes is enriched for functional teM genes, though still making up only a small fraction. For the vast majority of genes, developmentally responsive repression is mediated not by DNA methylation, but by polycomb related mechanisms (Baile *et al*. 2022).

Based on our findings, we suggest flagging genes that as having TE-like methylation. Based on this information, they can be given lower priority when identifying gene candidates for functional studies or omitted from genomics studies where their inclusion would introduce too much noise, not just in maize but in any plant genome. Annotation tracks showing teM genes from our study are now available on the NAM founder genome browsers hosted by MaizeGDB to make this information easily accessible. We also provide the IDs of all teM genes in all 26 genomes. Gene annotation remains a difficult and error prone task. Not alone, but along with other data such as synteny, expression, and known gene structural features, TE-like methylation can provide a valuable means of correcting faulty annotations.

### Gene body methylation is associated with stable gene copy number

We found that gbM genes were strongly underrepresented among tandem duplications (Fig. 3D). Since gbM genes tend to have stable expression over a broad range of cell types (Fig. 2E, F), we speculate that increased gene dosage of gbM genes tends to be more disruptive than that of UM genes. This would be consistent with prior work showing stronger genetic conservation of gbM genes across species (Takuno and Gaut 2013; Niederhuth *et al*. 2016; Seymour and Gaut 2020). Perturbing conserved processes that function in most cells is more likely to harm the plant than dispensable ones that function in a minority of cells.

### Gene body methylation epialleles are associated with weak but significant increases in expression

The maize NAM founder genomes, methylomes, and transcriptomes provide a powerful resource for studying relationships between methylation and gene expression. The quality of the genome assemblies and associated annotation allowed us to filter out genes whose CDS had diverged too much for useful comparisons, to accurately call methylation epiallele states in each genome; and to accurately access gene expression levels in each genome. We found that gbM epialleles correlate with expression increases relative to UM epialleles (Fig. 4A,B), indicating that Arabidopsis is not an anomaly in this regard (Shahzad *et al*.; Schmitz *et al*. 2013; Dubin *et al*. 2015; Meng *et al*. 2016; Muyle *et al*. 2021). The fact that the expression changes were small, with gbM genes showing ∼3% higher expression than UM genes, is consistent with the fact that gene body methylation is dispensable in some plants (Bewick *et al*. 2016). Maize, with its relatively high levels of genetic redundancy (Woodhouse *et al*. 2010) may be more tolerant of epiallele changes in individual genes than species such as Arabidopsis with more streamlined genomes.

### On the theoretical importance of gene body methylation

While the fact that similar studies in a dicot and a monocot species yield similar results points toward biological significance, it does not indicate that gbM is the causal factor in the correlation. It is conceivable–in fact, almost certainly true in some cases–that changes in gbM are a consequence of other factors that also affect expression. That gbM itself can positively affect gene expression is supported by recent unpublished data that genes that lose gbM in an Arabidopsis mutant also have reduced expression (Shahzad *et al*.). One possibility for how gbM could affect transcription is that it prevents internal transcription initiation that would interfere with normal transcription, as has been reported in Arabidopsis and in animals (Neri *et al*. 2017; Teissandier and Bourc’his 2017; Choi *et al*. 2020). However, DNA methylation and other linked chromatin modifications affect more than transcription. In theory, any process that involves enzymes making contact with DNA could be inhibited or facilitated (e.g., DNA repair, recombination, replication, transposon integration).

## METHODS

### Categorizing genes by methylation epiallele and metagene methylation analysis

CGmap files produced in the NAM founder study were used as the input for all DNA methylomes analyses (Guo *et al*. 2013; Hufford *et al*. 2021). The source Enzymatic methyl-seq reads are available at ENA ArrayExpress E-MTAB-10088. Each methylome was analyzed relative to its own reference genome, as opposed to the simpler but less accurate method of using B73 as the reference for all. The CGmapTools v0.1.2 mtr tool was used to calculate average methylation values of each gene using the “by region” method after filtering the CGmaps specifically for CDS using the CGmaptools select region tool (Guo *et al*. 2013, 2018). Version 1.0 gene annotations produced by the NAM assembly project were obtained from https://download.maizegdb.org. Only canonical gene annotations were used in defining CDS, as well as for all other genic features. Only genes with at least 40 cytosines in the CG context and 40 cytosines in the CHG context spanned by EM-seq reads were assigned methylation epialleles. UM epialleles were defined by both mCG and mCHG less than 0.05, gbM epialleles by mCG higher than 0.2 and mCHG less than 0.05, and teM epialleles by both mCG and mCHG methylation levels higher than 0.4. The metagene methylation values over 3 Kb upstream and downstream of genes and 1.5 Kb within genes were produced using the CGmapTools mfg tools with 100 bp intervals and minimum coverage of 1 (-c 1 parameter). First, however, the CGmapTools bed2fragreg tool was used in combination with an awk command to convert input gene annotations from BED format to the fragreg format used as input for the mfg tool.

### Identification of core genes and quantification of genic structural features

Gene IDs for core genes are provided at https://github.com/dawelab/Natural-methylation-epialleles-correlate-with-gene-expression-in-maize/tree/main/patterns%20of%20gene%20methylation/pangene_class. To identify core genes, which were annotated as genes in B73 and in all other 25 NAM founders, we made use of the pangene table that lists every gene in all 26 genomes with each row corresponding to a single pangene and each column as a single NAM founder (Hufford *et al*. 2021). The pangene table was downloaded from https://de.cyverse.org/anon-files//iplant/home/shared/NAM/NAM_genome_and_annotation_Jan2021_release/SUPPLEMEN TAL_DATA/pangene-files/pan_gene_matrix_v3_cyverse.csv. Tandem duplicate genes are included as multiple genes within a single cell. “NA” indicates a missing gene. Genome coordinates instead of a gene name indicate presence of homologous DNA but insufficient evidence for a gene annotation. These loci were not included in our analyses because they lack gene annotations, but corresponding genes in other genomes were still counted as core genes.

To calculate the length of genic structural features, we used awk commands using start and end coordinates in annotation files, and we summed the individual lengths per gene using the R aggregate(length∼gene,data,FUN=sum) function or summarise(sum(length)) function in gene unit. Intronic repeat lengths were obtained using the BEDTools v.2.30 (Quinlan and Hall 2010) intersect tool with -wo -wa -a (introns) -b (repeats). Zm-B73-REFERENCE-NAM-5.0.TE.gff3 repeat annotations are from https://download.maizegdb.org. The intersected repeat annotations were merged using the BEDTools merge tool to prevent overlapping repeat annotations from being counted twice. Since the gene annotation files in gff3 format lack intron annotations, an intron file for B73 was obtained using GenomeTools v.1.6.1 “-addintrons -retainids -sortlines” command. Exon coordinates were obtained from B73 annotation using the awk command to select the gene ID and exon column. Core genes were selected from the complete set using merge(coregene,input,all.x=T) in R, and the exon count for each obtained using aggregate(data,gene∼exon,FUN=length) in R.

To calculate numbers of genes with TE insertions, we used the Python pandas package (Python-3.7.4 environment) to generate simplified bed files for TE superfamilies. The BEDTools intersect tool with -wa -wb -a (intron) -b (TE) was used to identify TEs in introns. The R unique function was used to count each gene only once even if multiple annotations of the same TE superfamily were present. The R merge function was used to associate each gene with an insertion with its methylation epiallele status. The R table function was then used to count numbers of genes with at least one insertion for each superfamily.

To calculate extents of overlap between CDS and repetitive elements, we used the BEDTools intersect tool with -wo -a (CDS) -b (repeats). The R aggregate function was used to sum the CDS-overlapping repeat lengths per gene. The R merge function was used to associate each gene with its methylation epiallele status. “NA” was replaced with 0 for genes with no overlap between CDSs and repeats. The R table function was then used to count numbers of genes with CDS-overlapping repeat lengths that were greater than 100 bp.

### Calculating gene TPM values and assigning expression categories

To quantify gene expression, we mapped mRNA-seq reads from the NAM assembly project using similar methods to the NAM assembly project (Hufford *et al*. 2021). RNA-seq reads are available at ENA ArrayExpress E-MTAB-8633 and E-MTAB-8628. Briefly, STAR v2.7.2 software (Dobin *et al*. 2013) was used to map reads to each of the 26 genomes assemblies and their reference gene annotations. Unlike the NAM assembly project, however, gene annotations were used to guide read mapping with the --sjdbGTFfile and --twopassMode Basic parameters. Prior to read mapping, Cufflinks v2.2.1 (Trapnell *et al*. 2010) was used to convert gff3 gene annotations into gft format. As in the NAM assembly project, transcripts per million value (TPM), were obtained based on read counts per genes from featureCounts software v.1.6.0 (Liao *et al*. 2014) using uniquely mapping reads only (default method). For tissues with two mRNA-seq replicates, both replicates were merged.

To define the gene expression categories, we compared TPM values across the ten tissues (leaf tip, leaf middle, leaf base, root, shoot, ear, anther, tassel, endosperm, and embryo). We defined tissue-specific expression as TPM >= 1 in at least one tissue and TPM < 1 in at least one tissue, constitutive expression as TPM >= 1 in all ten tissues, and silent as TPM < 1 in all ten tissues. TPM values from all ten tissues were combined into one matrix, and the matrixStats package in R was used to identify each expression category.

### Identification of genes with syntenic homologs in sorghum

A table (NAM_subgenomes_vs_Sorghum_from_Hufford-2021.txt) linking maize genes to the two maize subgenomes and to their syntenic sorghum homologs was obtained from MaizeGDB (https://ars-usda.app.box.com/v/maizegdb-public/file/1091055382617). It was produced by Margaret Woodhouse using data from the NAM Founders Project and updated Dec 14, 2022. More details on its derivation are found in the same link and in the NAM Founders publication (Hufford *et al*. 2021). The R merge function (by=“gene”, all = T) was used to combine sorghum synteny information with other B73 gene information into a single table, and the R unique function was used to remove duplicated rows. Core genes were selected using the R filter function from the tidyverse package, and the numbers of genes with syntenic sorghum homologs were counted using the R table function.

### Conceptual summary of pangenes analysis

A schematic summary of our pangene methods is shown in Supplementary Figure 5. In brief, the three major input data sources—DNA methylomes, RNA transcriptomes, and gene annotations–were used to create a series of gene matrices, where the row coordinate indicates the pangene and the column coordinate indicates the NAM founder. The matrices included values such as CDS length, TPM, and epiallele status. Tandem duplicate genes were represented by lists of values within single cells. The initial matrices were intersected with each other using specific criteria such as CDS length to produce filtered matrices representing subsets of genes of particular interest. Values in these filtered matrices were then extracted and used as inputs for plotting distributions (such as TPM) or numbers of genes with specific features like stable epialleles.

### Identification of genes with intact CDS

For all pangene analyses, to exclude both annotation artifacts and genes with large structural changes in their CDSs, we required that the CDS length vary by less than 10% from the median of each pangene. Only singletons were included in determining the median length. Since CDS lengths for tandem duplicates were represented by lists of values within individual cells, the R rowMedians function was used to determine median lengths (rowMedians ignores values in lists). Both singletons and tandem duplicates were compared to the same singleton-defined median length and unqualified genes were removed using R logical operators. Removing genes by this criterion did not affect their core gene status, only whether they were included in subsequent analyses. Input gene sets for all pangene analyses were limited by this CDS length restriction.

### Identification of 1-to-1 and 1-to-N pangenes and counting epialleles

To identify pangenes with and without tandem duplicates, we created a pangene matrix of tandem duplicate counts, where a value of one indicated a singleton, a value of two indicated two copies, etc. The R rowMins and rowMaxs function were used to identify a set of pangenes with both singletons and duplicates. The R rowSums function was used to count numbers of singleton genes of each defined epiallele type (UM, gbM and teM) for each pangene. Pangenes that had more than one singleton epiallele type or had only one singleton gene with a defined epiallele were removed to produce the final 1-to-N pangene matrix. To count epialleles among tandem duplicates, the singleton epialleles (any cells not containing lists) were first converted to nulls. Then all cells containing lists were combined with the R unlist function and the epialleles counted with the R table function.

The stable 1-to-1 pangenes matrix was produced similarly as the 1-to-N pangenes, except it was derived from singleton-only pangenes. The unstable 1-to-1 pangene matrix was also derived from singleton-only pangenes, but only included pangenes represented by at least two epiallele types, where each epiallele type was represented by at least two genes. This produced four types of unstable 1-to-1 pangenes: UM-gbM, UM-teM, gbM-teM, and UM-gbM-teM.

### Comparison of epiallele expression levels in unstable pangenes

To compare epiallele expression values among unstable 1-to-1 pangenes, we calculated the mean TPM value for each epiallele type in each pangene individually using the following formula:

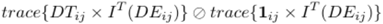

Variables are as follows: i is the pangene index and j is the genome index. DT_ij_ (tissue) is a matrix of TPM values, one for each tissue. DE_ij_ is a matrix of epialleles. I(DE_ij_) is an indicator matrix of zeros and ones, one indicator matrix for each of the three epiallele types. DT_ij_ (tissue) times the transpose of I(DE_ij_) and divide sum of the transpose of I(DE_ij_) over j was used to create three pangene lists for each tissue, where each list contains the mean TPM value for one 1epiallele type for that pangene. These lists were then used to calculate TPM differences for each epiallele type. To measure mean expression values of epiallele types in unstable 1-to-1 pangenes across the whole set, the R unlist function was used to make lists of TPM values for each epiallele type using the TPM matrices and epiallele matrix as inputs. The R summary function was used to calculate summary statistics from these lists. Pangenes had to have at least two genomes with each epiallele to be included in the unstable 1-to-1 pangenes set.

### mCG and TPM correlation coefficient calculations

To test for correlations between mCG and gene expression levels independently of epiallele category, we generated a pangene matrix of mCG values. Values for low confidence genes were replaced with “NA”. Low confidence was defined in the same way as other analyses: either CDS length varying by greater than 10% from the median of each pangene, or less than 40 cytosines in the CG context and 40 cytosines in the CHG context spanned by EM-seq reads. While there was no constraint on mCG, genes with mCHG values greater than 0.05 were excluded to avoid confounding effects of TE-like methylation. In addition to enrich for pangenes with variable mCG, the differences in mCG between the most extreme two genes of each pangene had to at least equal 0.2. The R functions rowMins(na.rm=T) and rowMaxs(na.rm=T) were used to calculate the minimum and maximum mCG values for each pangene. For the TPM matrices, positions matching ‘NA’ in the mCG matrix were also replaced with ‘NA’. Pearson correlations were calculated for each pangene using the R function cor.text(). The correlation coefficient was set to ‘NA’ if the standard variation of a pangene was 0 or if there were less than three available genes in a pangene to be tested. The mean and median of the correlation coefficients were calculated by the R summary() function.

### Other P-value calculations

To test whether the proportions of epiallele types differed between tandem duplicates of 1-to-N pangenes with minimum of two copies and 1-to-N pangenes with a minimum of four copies in Figure 3B and C, we applied Chi-square tests on epiallele counts using the R function chisq.test (x,y,correct=F). To test whether the direction of TPM differences between epiallele types was significantly different (gain or loss of expression in Figure 4B-D), we applied both binomial sign tests and Wilcoxon signed rank tests on each set of pangene counts. Binomial sign tests were done using the R function binom.test(sum(gbM>UM),n,p=0.5,alternative = “two.sided”) and the Wilcoxon signed rank test using the R function wilcox.test().

## Data Availability

Scripts and R methods used in this study are available at https://github.com/dawelab/Naturalmethylation-epialleles-correlate-with-gene-expression-in-maize

