## Supplemental Figures for "Natural methylation epialleles correlate with gene expression in maize"

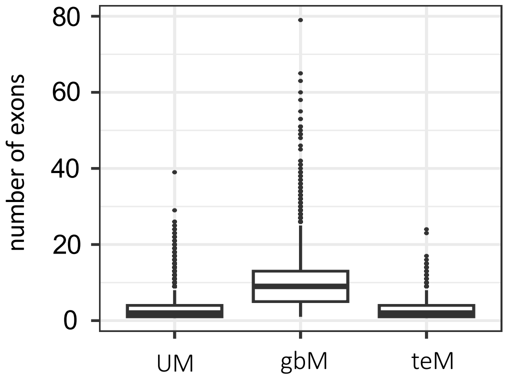


**Supplemental Figure 1: Exon counts**
Distributions of number of exons per gene in UM, gbM and teM genes (left to right). Differences between UM, gbM and teM genes were significant for all features (P-value < 10-10, Wilcoxon rank sum text).

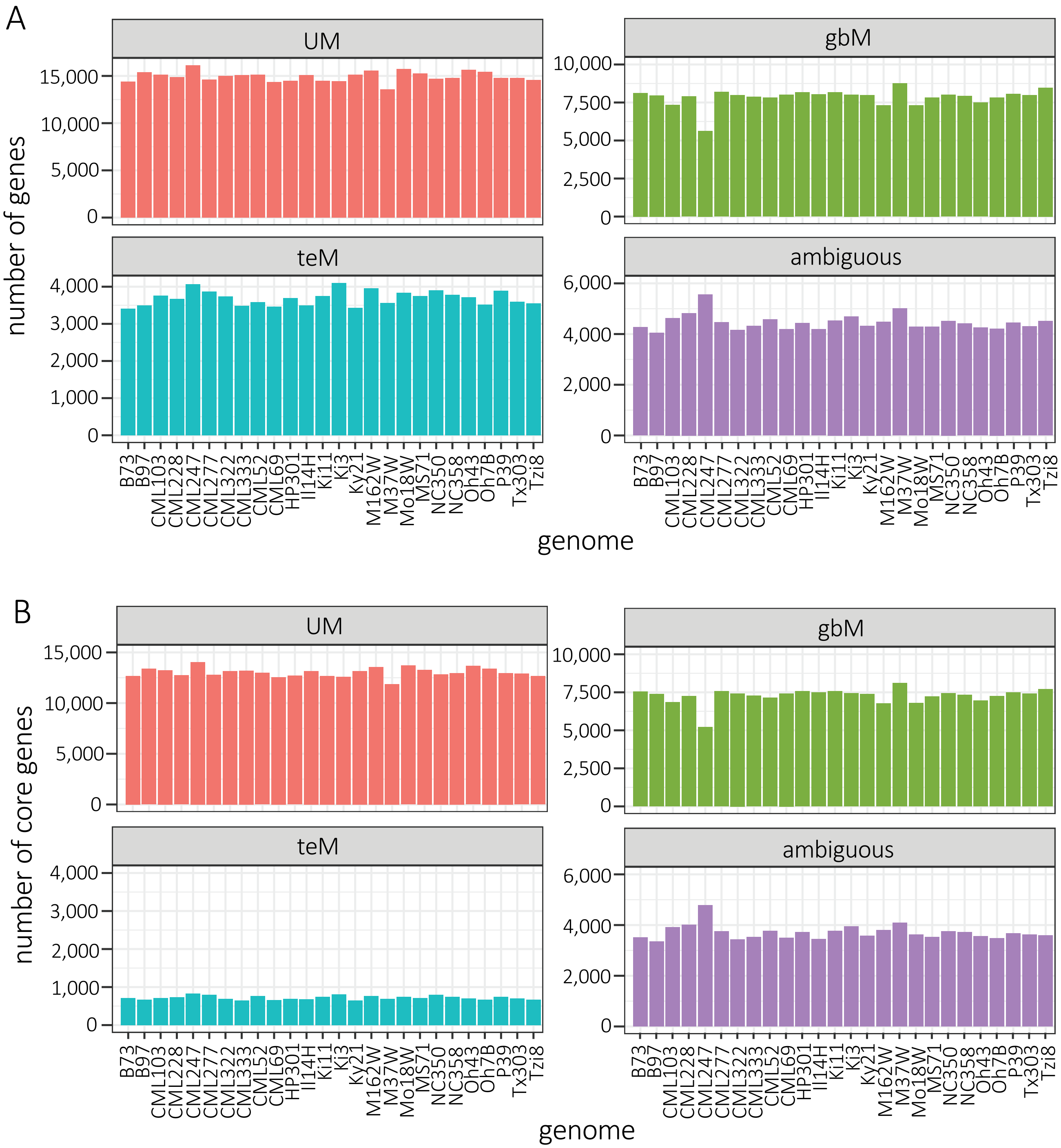


**Supplemental Figure 2. Numbers of genes with defined epialleles in NAM founder genomes**
**A.** Column heights indicate number of genes for UM, gbM, teM, and ambiguous genes for each NAM founder. Ambiguous genes only include ones with EM-seq read coverage of at least 40 CGs and 40 CHGs.
**B.** As in A, but only including core genes.

A

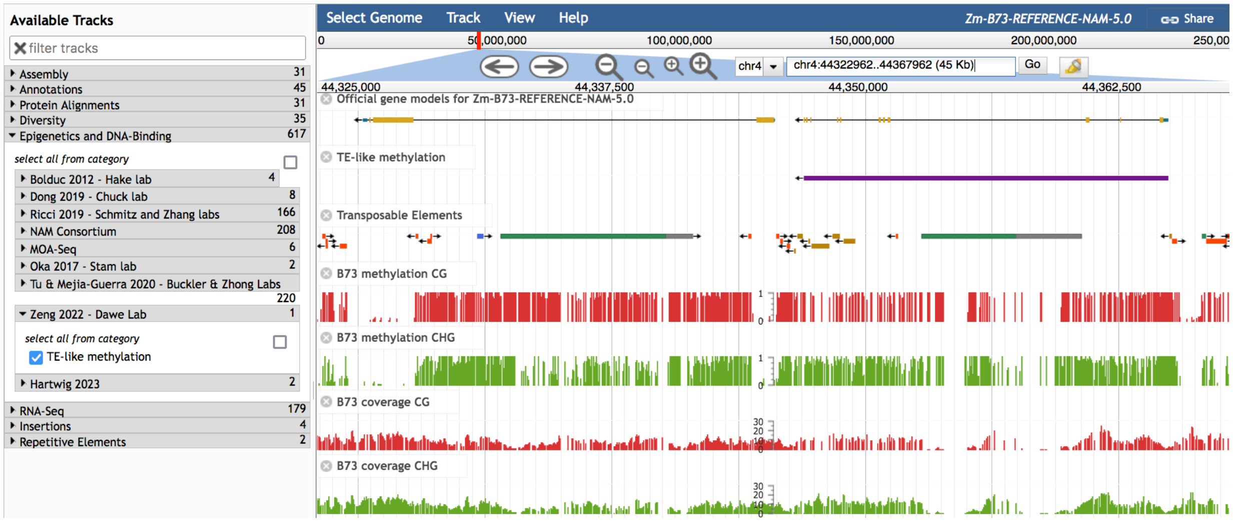


B

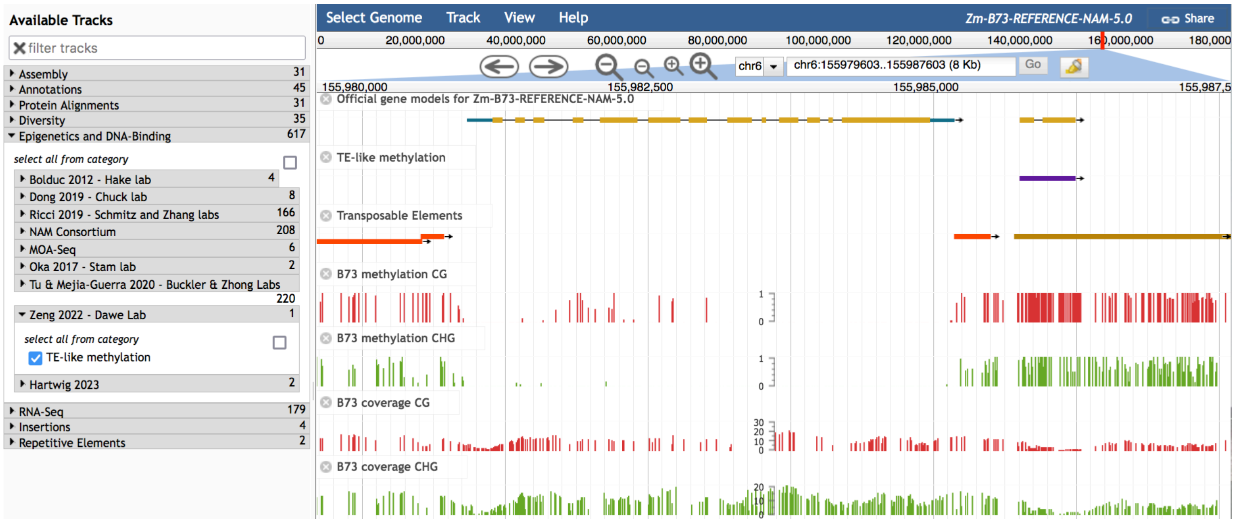


**Supplemental Figure 3: teM genes genome browser tracks**

**A.** JBrowse display of a 45-Kb region of the B73 v5 genome. The gene annotation on the left, Zm00001eb174770, is a UM gene as evidenced by high read coverage and low mCHG in exons. The gene annotation on the right, Zm00001eb174780, is a teM gene as evidenced by high read coverage and both high mCG and high mCHG in exons. The "TE-like methylation" track (purple) is available for all 26 NAM founder genomes. See the "Epigenetics and DNA-Binding" track options at https://jbrowse.maizegdb.org to find these tracks.

**B.** JBrowse display of an 8-Kb region of the B73 genome hosted by MaizeGDB, as in A. The gene annotation on the left, Zm00001eb287500, is an ambiguous gene with high read coverage but intermediate levels of mCG in exons. The gene annotation on the right, Zm00001eb287510, is a teM gene as evidenced by high read coverage and both high mCG and high mCHG in exons.

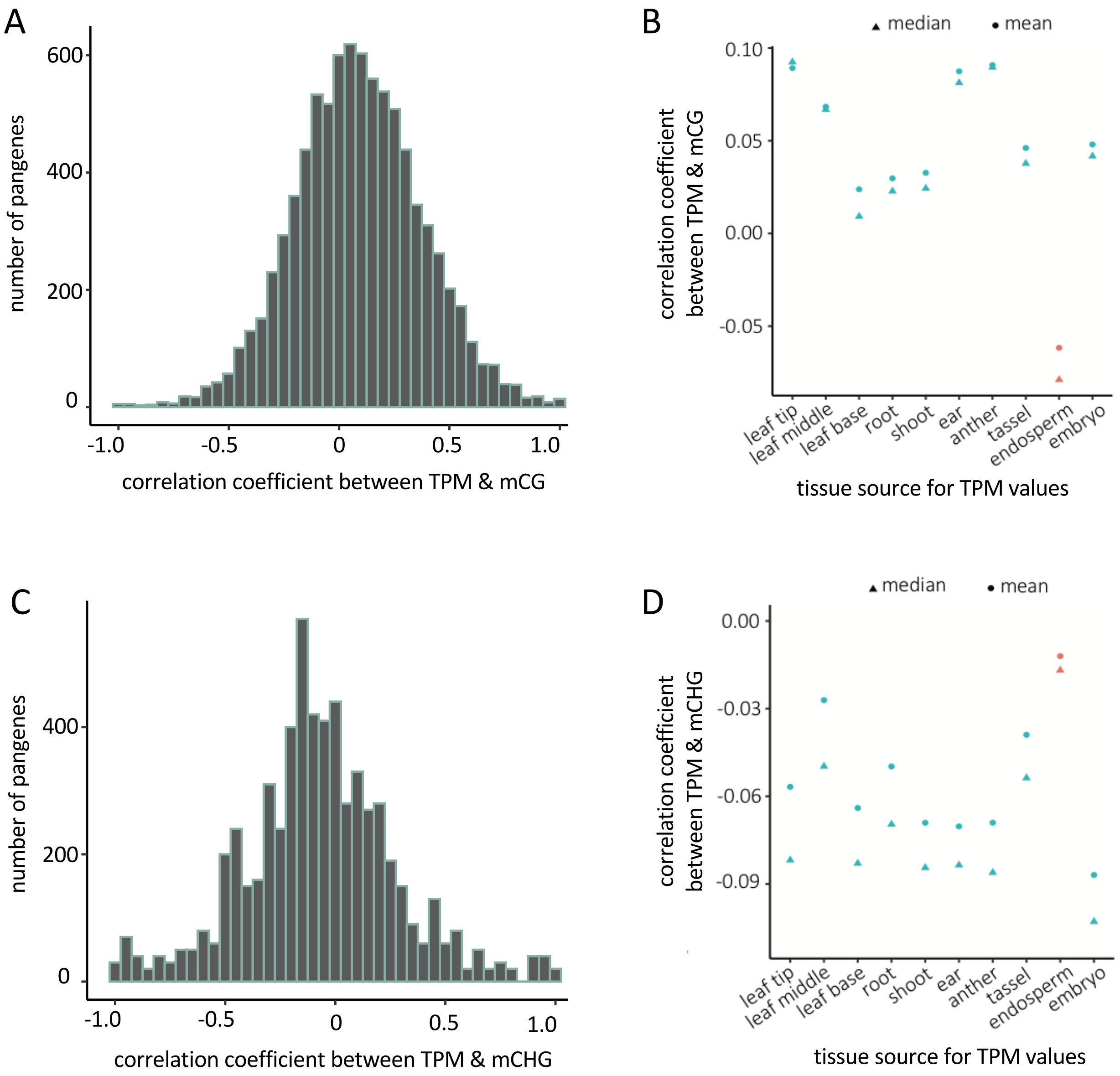


**Supplemental Figure 4: Correlations between methylation and gene expression**

**A.** Distribution of Pearson's correlation coefficients between mCG and TPM. Each pangene yielded one Pearson's correlation coefficient value based on mCG and TPM values for a minimum of three and maximum of 26 genes. TPM values are from ear tissue.

**B.** Mean and median Pearson's correlation coefficients for all pangenes. Each of the ten tissues used to calculate TPM values is shown separately.

**C.** Distribution of Pearson's correlation coefficients between mCHG and TPM. Each pangene yielded one Pearson's correlation coefficient value based on mCHG and TPM values for a minimum of three and maximum of 26 genes. TPM values are from ear tissue.

**D.** Mean and median Pearson's correlation coefficients for all pangenes. Each of the ten tissues used to calculate TPM values is shown separately.

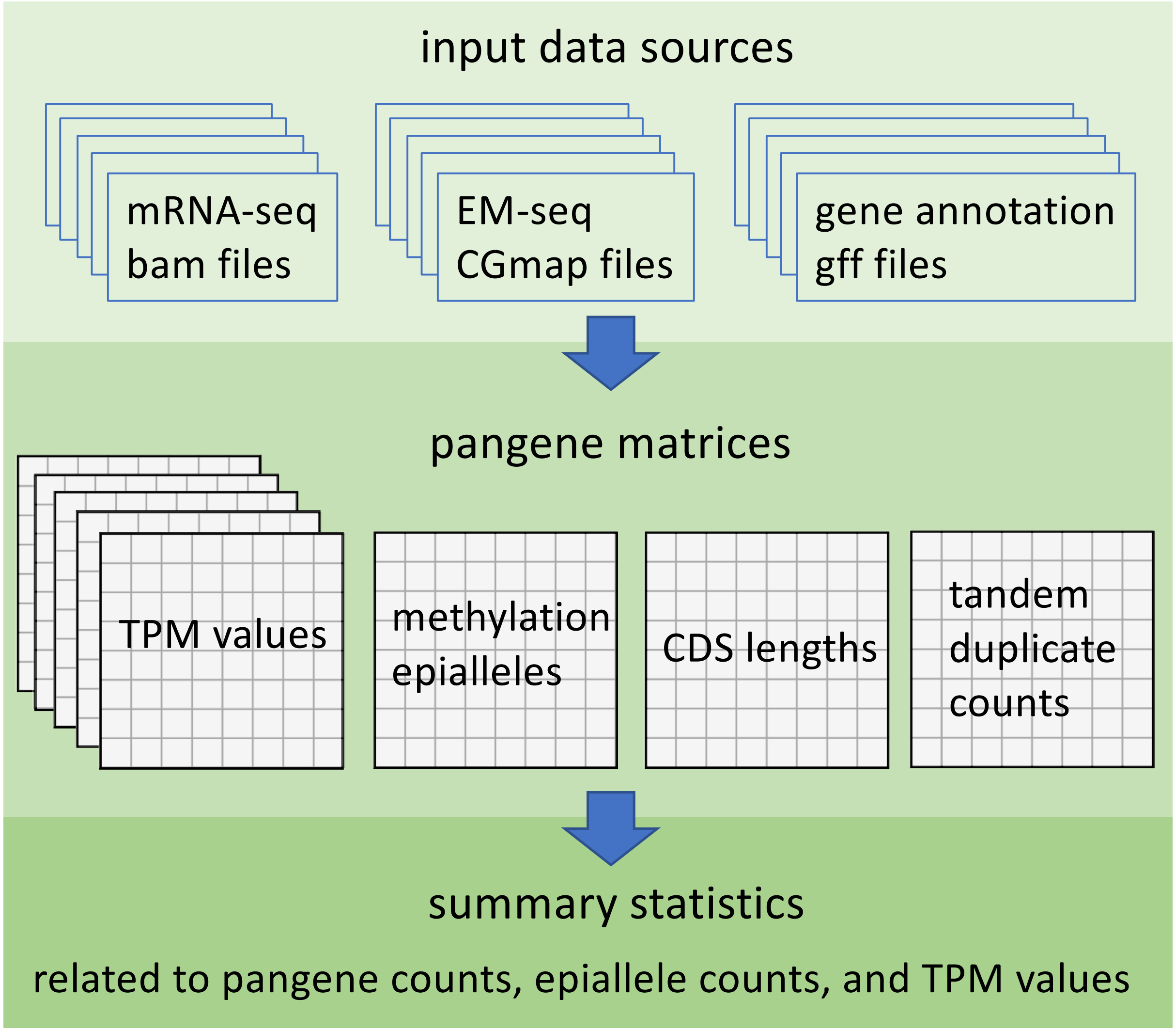


**Supplemental Figure 5: Schematic of pangene analysis workflow**
Input data sources for each NAM founder genome were either downloaded from maizeGDB or produced according to methods of the NAM assembly project (Hufford et al 2021). Pangene matrices consisted of one pangene per row. Each column corresponded to a NAM founder, with values from either single genes or from clusters of tandem duplicate genes.
